# JADE: Joint Alignment and Deep Embedding for Multi-Slice Spatial Transcriptomics

**DOI:** 10.1101/2025.11.21.689823

**Authors:** Yuanchuan Guo, Jun S. Liu, Huimin Cheng, Ying Ma

## Abstract

As spatially resolved transcriptomics (SRT) datasets increasingly span multiple adjacent or replicated slices, effective joint analysis across slices is needed to reconstruct tissue structures and identify consistent spatial gene expression patterns. This requires resolving spatial correspondences between slices while capturing shared transcriptomic features, two tasks that are typically addressed in isolation. Multi-slice analysis remains challenging due to physical distortions, technical variability, and batch effects. To address these challenges, we introduce Joint Alignment and Deep Embedding for multi-slice SRT (JADE), a unified computational framework that simultaneously learns spatial location-wise alignments and shared low-dimensional embeddings across tissue slices. Unlike existing methods, JADE adopts a roundtrip framework in which each iteration alternates between alignment and embedding refinement. To infer alignment, we employ attention mechanisms that dynamically assess and weight the importance of different embedding dimensions, allowing the model to focus on the most alignment-relevant features while suppressing noise. To the best of our knowledge, JADE is the first method that jointly optimizes alignment and representation learning in a shared latent space, enabling robust multi-slice integration. We demonstrate that JADE outperforms existing alignment and embedding methods across multiple evaluation metrics in the 10x Visium human dorsolateral prefrontal cortex (DLPFC) and Stereo-seq axolotl brain datasets. By bridging spatial alignment and feature integration, JADE provides a scalable and accurate solution for cross-slice analysis of SRT data.

## 1 Introduction

Spatially resolved transcriptomics (SRT) technologies provide high-throughput measurements of gene expression within tissue sections while preserving spatial context [6, 43, 47, 60, 28]. By enabling the joint study of molecular states and tissue architecture, SRT has transformed our understanding of developmental processes [9, 70], disease microenvironments [46, 81], and spatial cellular organization across diverse biological systems, from cancer [55, 5, 59, 26, 46, 1, 16] to neuroscience [42, 44, 52, 75]. As the resolution and scale of SRT continue to improve, there is growing interest in multi-slice SRT datasets, where multiple adjacent or replicated tissue sections are profiled to reconstruct 3D structures, map spatial trajectories, or assess reproducibility across individuals and conditions [19, 56, 21, 41, 14, 30, 40, 34]. However, analyzing multi-slice SRT data introduces a set of unique challenges. Each tissue slice may undergo non-linear spatial deformation during sectioning and mounting, while also exhibiting substantial variation in transcript capture efficiency and local tissue composition [77, 36]. These factors confound the alignment of corresponding regions across slices and obscure shared biological signals. Effective multi-slice integration must therefore address two tightly coupled tasks: spatial alignment and representation learning [38, 66, 35]. Alignment resolves spatial location -level correspondences across slices, while representation learning compresses high-dimensional gene expression data into a shared latent space that supports robust downstream analysis. Despite their interconnected nature, existing methods typically address these tasks in isolation, limiting their ability to perform coherent, biologically meaningful integration.

Previous approaches fall into two broad categories. Alignment-based methods [36, 73, 12, 27, 20], such as PASTE [77], estimate spatial location - level mappings across adjacent slices by jointly considering spatial coordinates and gene expression similarity, enabling reconstruction of 3D tissue volumes. However, these methods operate directly on raw expression profiles, which are often sparse, noisy, and affected by batch effects. These batch effects refer to systematic technical variations introduced during sample processing or sequencing, which can obscure true biological signals. As a result, they do not provide low-dimensional representations that are essential to downstream tasks such as spatial domain detection or trajectory inference. Conversely, representation learning–based methods [22, 33, 80, 78, 51, 23, 7, 72, 76], such as GraphST [37] and STAGATE [15], leverage graph neural networks to extract informative low-dimensional embeddings. While effective for extracting latent embeddings and identifying spatial domains, these methods typically process slices independently or jointly without explicitly resolving anatomical correspondences. As a result, homologous tissue regions may be represented inconsistently across slices, undermining interpretability and cross-slice comparison. To address the need for joint analysis across samples, integration methods originally developed for single-cell RNA sequencing data [71, 18], including Harmony [32], Seurat [56], and scVI [39], do not account for spatial context and assume shared coordinate systems across samples, assumptions that rarely hold in spatial data. Recently, approaches such as STAligner [79] and PRE-CAST [35] have extended integration techniques to spatial transcriptomics, but they either require additional input (e.g., batch id or histology image) or do not explicitly model spatial location-level alignment. These methods focus on harmonizing latent features across slices but cannot account for physical tissue distortions that are critical for spatial reconstruction.

Together, these limitations highlight the need for a unified framework that can resolve spatial correspondences and learn biologically meaningful representations across multiple slices, allowing alignment to guide representation learning, and vice versa. To address this need, we introduce JADE (Joint Alignment and Deep Embedding), a computational framework that integrates multi-slice SRT data by simultaneously learning (1) a probabilistic alignment between spatial locations and (2) a shared low-dimensional embedding space. JADE performs alignment in the latent space via attention-based optimal transport and enforces spatial and transcriptomic consistency through graph-based contrastive learning. This coupling ensures that learned embeddings are mutually aligned and biologically coherent, while correspondences between spatial locations across tissue slices respect both spatial geometry and gene expression structure.

To the best of our knowledge, JADE is the first method to jointly optimize spatial location-level alignment and representation learning within a unified, spatially informed model. We benchmark JADE on multi-slice SRT datasets from the human dorsolateral prefrontal cortex (DLPFC) [44] and the regenerating axolotl brain [67], and show that it consistently outperforms state-of-the-art alignment and embedding methods in spatial clustering accuracy, alignment fidelity, and biological interpretability. JADE offers a scalable and robust solution for integrative spatial analysis, particularly in settings that require the joint resolution of anatomical correspondence and functional representation across complex tissue landscapes.

## 2 Methods

### Problem Definition

Before we present our method JADE, we first formally introduce the problem setup. Consider a pair of SRT slices (*S*_1_, *X*_1_) and (*S*_2_, *X*_2_), where 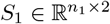and 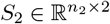 represent the spatial coordinates of *n*_1_ and *n*_2_ spatial locations in the two tissue slices, and 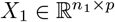 and 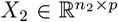 correspond to their respective gene expression matrices, where *p* represents the same set of genes measured across tissue slices. Given both gene expression and spatial coordinates for the two slices, our objective is to jointly learn: (1) an alignment matrix 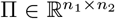 between these spatial locations, where Π_*ij*_ encodes the correspondence between spatial location *i* in slice 1 and spatial location *j* in slice 2, and (2) the low-dimensional embedding representations *H*_1_ and *H*_2_.

### Overview of JADE

Figure 1 illustrates the workflow of JADE. The pipeline begins with the input of SRT data from two slices (left panel). These inputs are passed through encoders to extract low-dimensional embeddings. Simultaneously, JADE infers cross-slice correspondences via a graph attention module to obtain embedding-space alignment. This module consists of a projection head, matrix multiplication, softmax, and iterative sinkhorn operations. To enhance generalizability, we employ a contrastive learning module that maximizes agreement between spatially neighboring locations while distinguishing dissimilar ones. A key innovation of JADE is its roundtrip learning scheme, in which embeddings and alignments are refined alternately within each training iteration. The outputs of JADE include a probabilistic alignment matrix between spatial locations across two tissue slices and embedding representation of each spot. These outputs support downstream analyses such as 3D tissue reconstruction (via the alignment matrix), spatial domain detection (via the learned embeddings), batch effect correction, and differential expression analysis. Detailed descriptions of each module are provided in the following sections and the pseudo code is shown in Appendix A. The implementation of JADE, along with data preprocessing scripts and pretrained models, is publicly available at https://github.com/YMa-lab/JADE.

**Figure 1.**
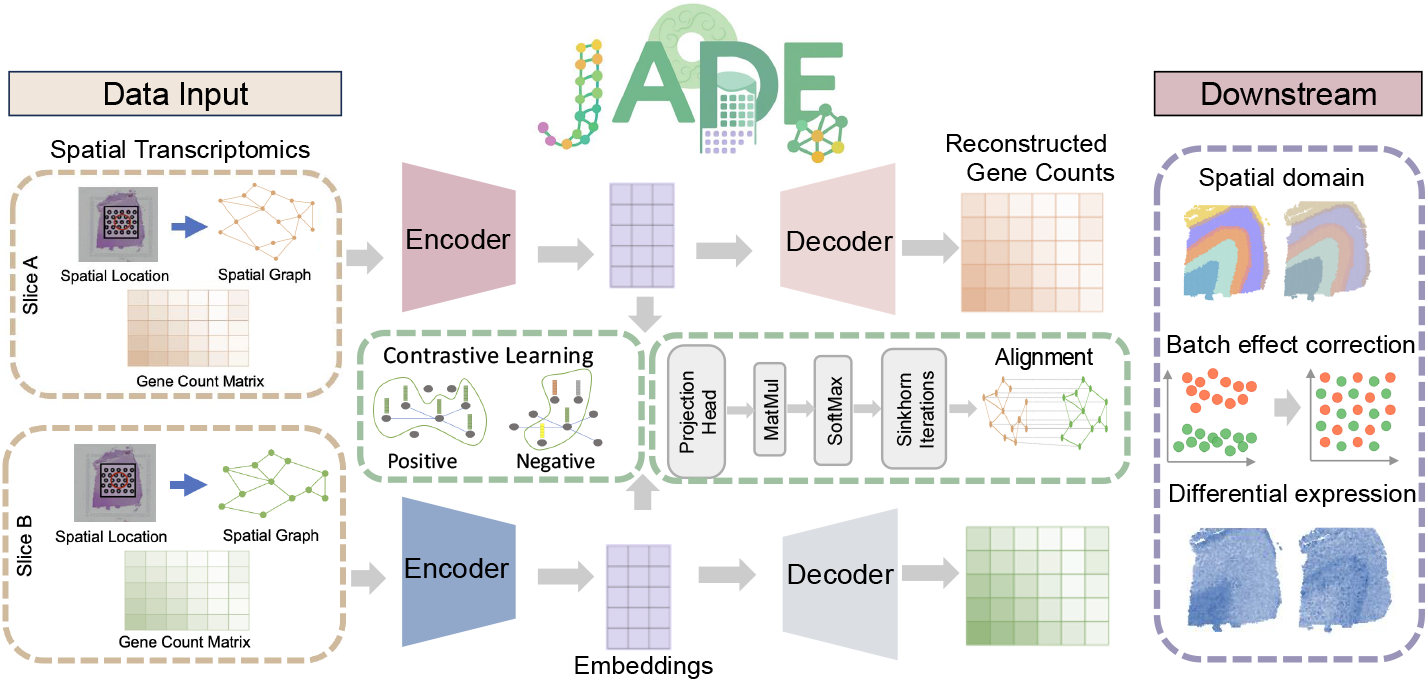
Workflow of JADE. Given two slices of SRT data (left box), JADE learns a probabilistic alignment and low-dimensional embeddings simultaneously. One unique feature of JADE is that embedding and alignment are updated in a roundtrip way in which each iteration alternates between alignment and embedding refinement. Downstream applications (right box) include spatial domain detection, batch effect correction, and differential expression analysis.

### 2.1. Graph Autoencoder

Let 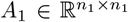, and 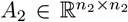 denote the two adjacency matrices of the spatial graph in two slices obtained using the method described in Appendix B. Given two SRT slices (*A*_1_, *X*_1_) and (*A*_2_, *X*_2_), we first encode them independently using graph convolutional networks (GCNs) [31] to obtain initial embeddings. Each encoder aggregates information from neighboring spatial locations to learn low-dimensional embeddings using a one-layer GCN: *H*_1_ = Relu(*Ã*^1^*X*_1_*W*_1*e*_ + *b*_1*e*_),*H*_2_ = Relu(*Ã* ^2^*X*_2_*W*_2*e*_ + *b*_2*e*_), where 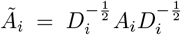 is the degree normalized adjacency matrix, ensuring proper feature scaling across neighboring locations. Without normalization, a node with more neighbors would accumulate disproportionately large feature values, leading to biased representations. *W*_*ie*_ and *b*_*ie*_ are trainable weight and bias parameters of the encoder for slice *i* ∈{1, 2} . ReLU is applied as a nonlinear activation function to enhance feature expressiveness. 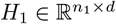 and 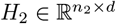 are the latent embeddings of two slices, where *d* is a hyperparameter that defines the latent dimension.

To ensure that the learned embeddings retain key biological information, we decode them back into the original feature space using a GCN-based decoder 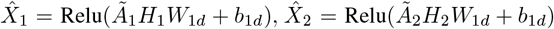, where *W*_*id*_ and *b*_*id*_ represent trainable parameters of the decoder for slice *i ∈* {1, 2}. The reconstruction loss ensures that the decoded outputs 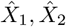 accurately recover the original gene expression profiles 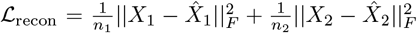, where *∥ · ∥* _*F*_ denotes the Frobenius norm, measuring the difference between the reconstructed and original feature matrices.

### 2.2. Graph Attention for Embedding-Space Alignment

Given embeddings *H*_1_, *H*_2_, we obtain the alignment matrix Π through the following steps.

#### Step 1: Compute cross-attention weights

We first compute an attention-based similarity matrix 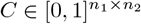to measure correspondences between embeddings of two slices:

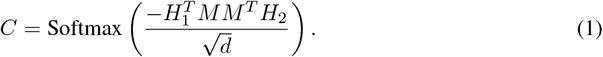

Here *M ∈* ℝ^*d×d*^ is a learnable linear projection implementing an attention mechanism [63], enabling the model to focus on key cross-slice features.

#### Step 2: Obtain the alignment matrix from cross-attention weights

The cross-attention weights *C* encode pairwise embedding similarities, but it is only row-stochastic: each row sums to 1, yet the column totals are unrestricted. This imbalance can lead to biased mappings, where some target locations accumulate disproportionately high mass, while others receive very little.

To achieve a balanced alignment, we reinterpret the alignment problem within an *optimal transport* (OT) [61] framework, enforcing both rows and columns to satisfy uniform marginal constraints. In this framework, the alignment matrix satisfy 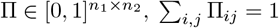. Each element Π_*ij*_ denotes the amount of probability mass transported from source point *i* to target point *j*, with the constraint that the entire matrix satisfies specific marginal distributions. Specifically, we apply the Sinkhorn-Knopp algorithm [54, 13] to transform the raw similarity scores in *C* into a doubly-stochastic alignment matrix Π. This iterative procedure alternately normalizes rows and columns, enforcing each row sum to 1*/n*_1_ and each column sum to *n*_2_, ensuring that every spatial location contributes and receives an equal amount of alignment mass. Thus, the alignment becomes fair and balanced, avoiding situations where a few locations dominate the mappings. Additionally, we introduce a marginal regularization term into our objective function: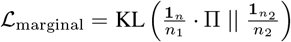. This penalty is defined as the KL divergence between the column sums of Π and the desired uniform marginal distribution. In doing so, we explicitly discourage deviations from uniformity, further reinforcing balanced alignments.

#### Step 3: Spatial and embedding aware alignment losses

To refine the alignment and maintain spatial structure, we define two primary losses motivated by the fused Gromov-Wasserstein distance [45, 64]: (1) Spatial structure preservation loss (Mis-Maintain Loss) ensures that spatial relationships in both slices remain consistent after alignment 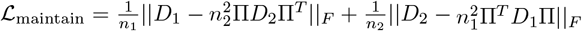, where *D*_1_ and *D*_2_ are pairwise spatial distance matrices. Each entry *d*_1*ij*_ (for slice 1) and *d*_2*ij*_ (for slice 2) represents the squared Euclidean distance between spatial locations *i* and *j*. The scaling factors 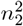 and 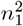 ensure that the transported distance matrices Π*D*_2_Π^*T*^ and Π^*T*^ *D*_1_Π remain on the same scale as the original *D*_1_ and *D*_2_, compensating for the normalization inherent in Π. Minimizing ℒ_maintain_ encourages spatial locations that were close to each other before alignment to remain close after alignment, while spatial locations that were far apart should continue to be far apart. (2) Embedding alignment loss (Mis-Alignment Loss) ensures that the aligned embeddings are close to each other, maintaining meaningful biological correspondence 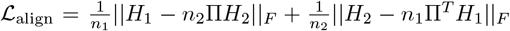 . Similarly, the scaling factors *n*_2_ and *n*_1_ensure that the transported embedding matrices *n*_2_Π*H*_2_ and *n*_1_Π^*T*^ *H*_1_ remain on the same scale as the original *H*_1_ and *H*_2_. The first term in the above equation minimizes the discrepancy between slice 1 embeddings and their aligned counterparts from slice 2. The second term enforces the same alignment constraint in the reverse direction.

### 2.3. Self-supervised Graph Contrastive Learning

We refine our embeddings via self-supervised contrastive learning similar to GraphST [37], using the Graph Infomax objective [65] to maximize mutual information between each spot and its local neighborhood. This drives adjacent, biologically similar spots closer in latent space while pushing structurally distinct or distant spots apart.

For each spatial location *i*, we define its local spatial proxy *r*^*i*^ as: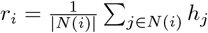, where *N* (*i*) denotes the set of immediate neighbors of spatial location *i, h*_*j*_ is the embedding of neighboring spatial location *j*. The pair (*h*_*i*_, *r*_*i*_) forms a positive pair, reflecting spatial proximity and biological similarity. To create negative examples, we generate a shuffled embedding matrix *H*^*′*^ by randomly permuting the rows of *H*, destroying spatial coherence. For each *i*, let 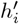 be the shuffled embedding and 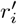 its corresponding shuffled proxy, forming a negative pair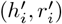. Finally, we train a discriminator Φ via binary cross-entropy loss to distinguish these positive from negative pairs. The slice-specific overall contrastive loss is: 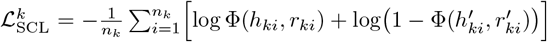 for *k* = 1, 2. The contrastive loss is: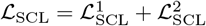 .

### 2.4. Final Loss Function

In summary, the training objective of JADE comprises three components:

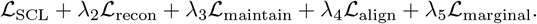

Throughout the paper, we set *λ*_2_ = 10, *λ*_4_ = 0.1, *λ*_5_ = 1. We use a data-driven method to select *λ*_3_ from 0.2 to 2.0 depending on a calculated similarity score between slices. If two slices are similar, we use a larger *λ*_3_ encourage information sharing; otherwise, we use a smaller *λ*_3_ to prevent negative transfer. A comprehensive description of hyperparameter tuning procedures is provided in Appendix E. Furthermore, the contribution of each individual component in JADE is evaluated through systematic ablation studies, as detailed in Appendix F. Results show that performance deteriorates markedly when any loss term is omitted, demonstrating that both the misalignment and mismaintain losses are crucial for optimal embedding and alignment quality.

#### Downstream analysis

After training, we obtained two sets of low-dimensional embeddings. We normalized the length of each embedding vector and applied the mclust algorithm independently, specifying the cluster number based on prior knowledge. Following [79, 37], we set the cluster number from 5 through 7 for the DLPFC dataset, and following [20], from 16 through 17 for the axolotl brain dataset, with each cluster corresponding to a distinct cell type. We then leveraged the inferred domains to conduct differential expression analysis. For the batch-effect correction evaluation, we projected the embeddings into a 2-dimensional space using UMAP, visualized their spatial distribution, and computed the local Simpson diversity index to quantify how well the embeddings from different slices are intermingled. To assess alignment quality, we visualized and quantified the highest-probability correspondences in the alignment matrix within each ground-truth domain, explicitly marking correct and incorrect matches, and computed accuracy scores to measure alignment performance.

## 3 Fast Computation for JADE

A major challenge for SRT alignment methods is their high computational cost, scaling quadratically with the number of spatial locations. Methods such as PASTE [77] and our JADE algorithm require computing similarity matrices between all location pairs, resulting in prohibitive runtime and memory usage for high-resolution datasets. The primary computational bottleneck in JADE is the crossattention step, which calculates pairwise similarities and scales as *O*(*n*_1_*n*_2_*d*). To accelerate this, we introduce an approach that aligns at a coarser resolution using aggregated spatial units (“hyperspots”), significantly reducing computational complexity. Despite this approximation, Fast-JADE achieves performance comparable to JADE on the DLPFC dataset (see Appendix G for detailed comparison).

### Hyperspot embedding construction

For each tissue slice, we apply *K*-means clustering to group spatial locations into fewer, coarser spatial units, resulting *m*_1_*≪ n*_1_ and *m*_2_*≪ n*_2_ hyperspots respectively. The number of hyperspots is set to approximately 10-20% of the original number of spatial locations for each slice. Each hyperspot embedding is computed by averaging embeddings of the spatial locations within the cluster, resulting in compact representations.

### Compute cross-attention between hyperspots

We compute the cross-attention weights and alignment matrix at the hyperspot level. Specifically,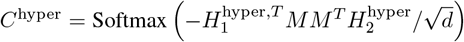 is similar to (1). Using these hyperspot-level attention weights, we then compute the alignment and maintenance losses, *ℒ*_maintain_ and *ℒ*_align_ at the hyperspot level.

### From hyperspot-level to spatial location-level Alignment

After training, we transfer the learned alignment back to the full-resolution space. Specifically, we reuse the same projection head *M* to compute the fine-grained spatial location-wise attention weights: 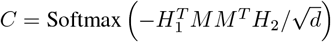,where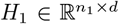 and 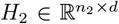 are the original spatial location embeddings. This ensures consistency between coarse and fine levels and avoids retraining at full resolution. The resulting matrix *C* is then passed through the Sinkhorn-Knopp algorithm to obtain an alignment matrix Π with uniform marginals.

## 4 Experiments

We benchmarked JADE against six representative baseline models, selected for their relevance to either spatial alignment or embedding tasks in spatial transcriptomics. For alignment accuracy, we compared with PASTE [77], Seurat [56], and STAligner [79], three widely used methods that align SRT slices based on transcriptomic similarity and spatial proximity. For evaluating the learned embeddings through spatial domain detection, we included GraphST [74], STAGATE [15], and STAligner, which incorporate spatial or graph-based information to enhance domain identification.

To provide a unified assessment, we used a comprehensive alignment metric that accounts for both correctly aligned and unaligned locations (Appendix B). Embedding quality was quantified using the Adjusted Rand Index (ARI) for biological domain recovery, the integrated local inverse Simpson’s index (iLISI) [53] for cross-slice integration, and UMAP visualizations for qualitative evaluation of coherence and batch correction. For biological interpretability, we conducted differential expression analysis to identify domain-specific marker genes and validated them against known anatomical annotations and literature, confirming that the learned embeddings capture biologically meaningful spatial structures and gene associations.

We also tested JADE on two additional datasets in Appendix H: the MERFISH dataset [10] and the Breast-Cancer Visium/Xenium dataset [25], previously used in SLAT [69]. As summarized in Table 11, JADE consistently outperforms or matches state-of-the-art baselines, including SLAT, STAligner and SpotScape[49], in both domain-detection accuracy (ARI) and alignment accuracy (ACC). These results demonstrate JADE’s robustness and adaptability across distinct spatial transcriptomics platforms (MERFISH, Visium, Xenium) and tissue types (brain and breast).

### 4.1. Joint Spatial Alignment and Representation Learning Across Human DLPFC Slices

#### Dataset description

We evaluated our proposed method on the Human Dorsolateral Prefrontal Cortex (DLPFC) dataset generated with the 10x Visium platform [55], a standard benchmark in spatial transcriptomics. The dataset comprises 12 serial tissue sections, including four sequential slices (A–D) from each of three donors (I–III), with expression profiles of 33,538 genes measured at 47,681 spatial spots. Within each individual, slices A-B and C-D are adjacent pairs separated by 10 *µ*m, while slices B and C have a larger separation of 300 *µ*m. Differences across individuals are primarily attributed to batch effects arising from technical variations in sample processing, sequencing, or tissue handling. Each slice was annotated with seven spatial domains, corresponding to six cortical layers and the white matter [44]. These expert-provided annotations served as the ground truth for spatial domain detection analysis. Following preprocessing in Appendix B, the two selected slices retained about 1500 shared highly variable genes for each pair.

#### Improved clustering accuracy

Figure 2(A) provides a visual comparison of the predicted spatial domains from each method against the ground truth for both Slice A and Slice B of Sample III. Visually, JADE demonstrates the highest accuracy in recovering the annotated cortical structure, across slices A and B with clear, smooth boundaries. In contrast, other methods yield noisier and less coherent spatial domains. Quantitatively, JADE achieves median ARI values of approximately 0.61 and 0.65 for Slices A and B, for Sample III as shown in Figure 2(B). This represents a statistically significant improvement over GraphST (0.59 and 0.58), STAligner (0.56 and 0.59), and STAGATE (0.52 and 0.53). The top row of Figure 2(C) further visualizes the learned low-dimensional embeddings produced by each method using UMAP, colored by their true biological cluster annotation. JADE consistently demonstrates superior performance, yielding highly distinct and well-separated biological layers that form cohesive clusters with minimal overlap. Appendix C shows additional results for other samples and other adjacent slices.

**Figure 2.**
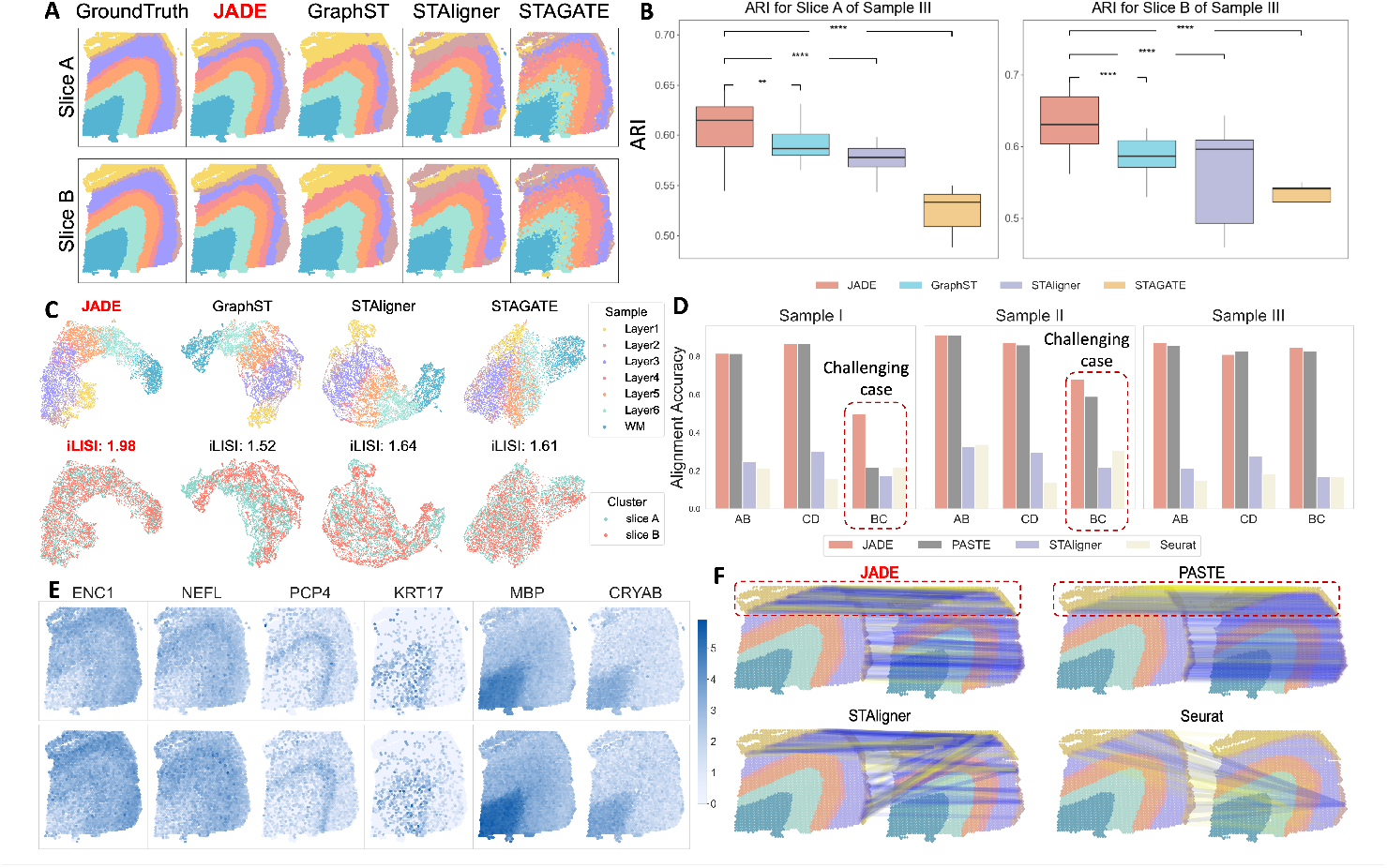
(A) Predicted spatial domains for slices A and B (Sample III) using five methods. (B) ARI for slices A and B of Sample III in DLPFC. *p*-values are calculated by Wilcoxon-rank sum test, where ** *p <* 1%,\*\*\**p <* 0.1% and **** *p <* 0.01% (C) UMAP of embeddings colored by predicted clusters (top) and slice (bottom). (D) Alignment accuracy across adjacent slice pairs (AB, CD, BC). (E) Spatial expression of marker genes confirms biologically relevant domain structures. (F) Visual comparison of alignment accuracy for Layer 1 in Sample III (Slice A vs. Slice B). True alignments are represented in blue; wrong ones are shown in yellow.

#### JADE effectively mitigates batch effects in multi-slice integration

The bottom row of Figure 2(C) illustrates the effectiveness of batch effect removal across different methods by visualizing UMAP embeddings colored by slice identity (slice A: blue, slice B: red). Effective integration is reflected by the intermixing of spatial locations from different slices, indicating successful alignment in a shared latent space. JADE achieves the highest inter-slice mixing, suggesting excellent batch correction performance. In contrast, GraphST, STAligner and STAGATE display limited alignment: the two slices remain largely separated within the UMAP space, suggesting poor batch removal efficacy. JADE also attained the highest iLISI score of 1.98, outperforming GraphST (1.52), STAligner (1.64), and STAGATE (1.61), confirming its superior performance in integrating SRT data across slices while correcting for batch effects.

#### Superior alignment performance

Figure 2(D) presents a comprehensive quantitative assessment of alignment accuracy across adjacent slice pairs (AB, BC, CD) for three distinct samples, comparing JADE against existing methods including PASTE, STAligner, and Seurat. Among these, the B–C pairs represent the most challenging alignment scenario due to their larger inter-slice distance (300*µm*),compared to the 10*µm* separation in A–B and C–D pairs. Across all samples and slice combinations,JADE consistently demonstrates superior alignment accuracy, particularly in challenging scenarios.While PASTE achieves comparable performance to JADE in easier settings (AB and CD pairs), it exhibits substantial performance degradation when confronted with the more challenging BC alignments. Notably, for BC pairs in samples I and II, JADE surpasses PASTE by margins exceeding 50%. STAligner shows moderate accuracy but is consistently lower than JADE and PASTE across all slice pairs. Seurat performs the poorest in all cases, with particularly low alignment accuracy in the BC pairs. These results highlight JADE’s superior alignment performance, particularly in challenging integration settings. In Figure 2(F), we visualize Layer 1 alignment between Slice A and Slice B in Sample III across four methods. Blue lines indicate correct matches within the same annotated layer, while yellow lines indicate errors. JADE achieves the best outcome, with mostly correct (blue) alignments and very few incorrect (yellow) ones.

#### Domain-specific differential expression gene analysis

We performed domain-specific differential expression (DE) analysis based on the spatial domains detected by JADE and identified six representtative marker genes (*ENC1, NEFL, PCP4, KRT17, MBP*, and *CRYAB*). These genes were selected for their strong domain-level enrichment and subsequently validated through literature for their well-established region-specific expression patterns in the human cortex. Figure 2(E) displays the spatial expression patterns of these genes across two adjacent DLPFC tissue slices (top and bottom rows). *ENC1, NEFL*, and *PCP4* exhibit clear laminar structures consistent with neuronal populations localized to middle and deep cortical layers [44, 68, 17]. Their spatial localization is preserved across slices, demonstrating coherent biological structure. *KRT17*, typically associated with epithelial-like or glial cells, shows more punctate and scattered expression, particularly in the upper portions of the slices [11, 44]. *MBP*, a myelination-related gene, is strongly expressed in the lower regions of both slices, likely corresponding to white matter (WM), and the spatial coherence across slices further supports successful alignment [48, 44, 24]. *CRYAB*, a stress-response and astrocyte-associated marker, exhibits a broader expression domain with intermediate intensity [4].

Beyond expression-based baselines, we further evaluated JADE against GPSA [27], an image-informed alignment model that integrates histological features in Appendix H. As summarized in Table 10, JADE achieves higher alignment accuracy across all DLPFC slice pairs while maintaining comparable clustering quality, indicating that reliable alignment can be achieved without reliance on histology images.

### 4.2. Joint Learning of Axolotl Brain Dataset Across Different Developmental Stages

We applied our method to the task of jointly learning alignments and embeddings of SRT data across distinct developmental stages. This integration is crucial for gaining insights into the dynamic processes of cell proliferation and differentiation. However, achieving such comprehensive integration is inherently challenging due to technical batch effects and fundamental biological shifts, both of which significantly complicate the establishment of accurate underlying alignment.

#### Dataset description

We evaluated JADE on a SRT dataset of the axolotl telencephalon (a region of the brain) generated using the Stereo-seq platform [9]. This dataset captures gene expression across five developmental stages: three embryonic stages, the juvenile stage, and the adult stage. In this study, we focused our analysis on the last two, which together span a broader range of mature brain architecture. The juvenile stage and adult stage have 17 and 16 distinct cell types respectively. Among them, fourteen are shared between both stages, while the rests from the juvenile stage differentiate or transition into new cell types in the adult stage. Following preprocessing, the two selected slices retained 1,000 shared highly variable genes, with 11,698 and 8,243 spatial locations, respectively. To reduce computational time, we applied the accelerated JADE method introduced in Section 3, using 1,000 hyperspots per slice, corresponding to about 10% of the spatial locations.

#### Improved clustering accuracy

Figure 3(A) provides a visual comparison of the domains detection results of JADE and baseline methods in Adult (top row) and Juvenile (bottom row) slices. Compared to the ground truth, JADE consistently recovers structurally coherent and anatomically accurate domains in both stages. Notably, in the adult brain, JADE is the only method that successfully reconstructs the blue peripheral ring cluster, representing vascular leptomeningeal cells (VLMC) [62]. This region is either partially fragmented (in GraphST), blurred and mixed with neighboring domains (in STAGATE, STAligner). In addition, JADE accurately produces a coherent, bilaterally symmetric red region in the telencephalon core, while STAGATE yields considerable color mixing within this region, STAligner only partially recovers it, and GraphST produces an overexpanded red region, suggestive of excessive smoothing. These qualitative patterns are supported by the quantitative results in Figure 3(B). For the Adult stage, JADE achieves the highest median ARI score (approximately 0.53), significantly outperforming STAligner (0.48), STAGATE (0.42), and GraphST (0.43). For the Juvenile stage, JADE also obtains a high median ARI (approximately 0.37), demonstrating statistically significant improvement over STAGATE (0.35) and GraphST (0.32). While JADE’s performance for the Juvenile stage is quantitatively comparable to STAligner’s by ARI, its qualitative domain recovery remains notably more coherent and anatomically accurate as shown in Figure 3(A). The top row of Figure 3(C) further visualizes the UMAP for the embedding features, colored by the true cluster annotation. JADE produces well-separated and compact clusters, indicating a clear differentiation of spatial domains in the latent space, while other methods exhibit considerable mixing of domain colors, reflecting ambiguity in domain boundaries.

**Figure 3.**
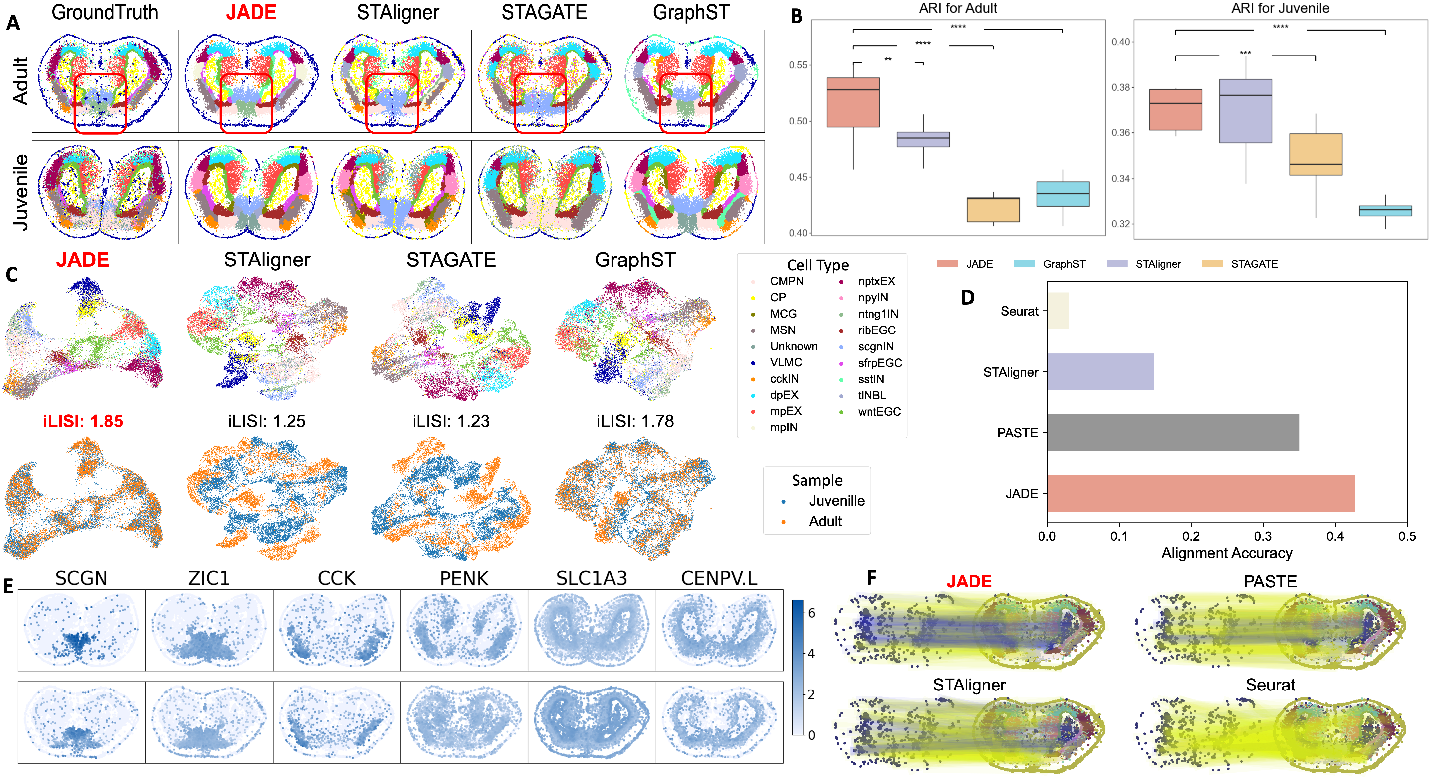
(A) Domain segmentation results for Juvenile and Adult slices comparing ground truth against four methods. (B) ARI scores for four methods. *p*-values are calculated the same as in Figure 2. (C) UMAP of embeddings colored by predicted clusters (top) and slice (bottom). (D) Alignment accuracy for Juvenile and Adult slices. (E) Spatial expression of marker genes confirms biologically relevant domain structures. (F) Visual comparison of alignment accuracy for CP cell type between the juvenile and the adult. True alignments are represented by blue lines, while wrong alignments are shown in yellow.

#### JADE effectively mitigates batch effects in multi-slice integration

The bottom row of Figure 3(C) compares methods via UMAP embeddings colored by slice (Adult: red, Juvenile: blue). JADE shows near-perfect mixing within structured clusters, indicating superior batch-effect removal. In contrast, STAligner, STAGATE, and GraphST display notable batch-induced separation. Quantitatively, JADE achieves the highest LISI score (1.85), surpassing GraphST (1.78), STAligner (1.25), and STAGATE (1.23), highlighting its effectiveness in integrating data across developmental stages.

#### Superior alignment performance

As shown in the bar plot in Figure 3(D), JADE achieves the highest alignment accuracy among all methods, substantially outperforming PASTE, STAligner, and Seurat. This result underscores JADE’s ability to establish precise correspondences between spatial locations across slices, even under the challenging setting of cross-developmental stage alignment, where substantial morphological and transcriptional variation exists. Figure 3(F) further visualizes the alignment accuracy for cortical plate cell type between the Juvenile slice and the Adult slice, to support this finding. JADE yields the highest proportion of correct (blue) alignments, demonstrating its superior ability to preserve anatomical consistency during cross-slice registration.

#### Domain-specific gene

Figure 3(E) displays the spatial expression patterns of six canonical marker genes, *SCGN, ZIC1, CCK, PENK, SLC1A3, CENPV*.*L*, across Juvenile and Adult slices (top and bottom rows). These genes are known to exhibit region-specific expression within the axolotl brain and thus serve as internal benchmarks for spatial alignment and biological interpretability. Specifically, *SCGN* and *ZIC1*, for example, are associated with distinct neuronal populations and developmental patterning, and their laminar or domain-restricted expression patterns are preserved across developmental stages, reflecting coherent spatial organization [2, 50, 58]. *CCK* and *PENK*, which are neuropeptide-related genes, show distinct regional enrichment, highlighting the emergence of functional specialization in the maturing brain [29, 8]. *SLC1A3*, a glutamate transporter gene involved in astrocytic function, exhibits broad but domain-enriched expression, consistent with known glial distribution [3]. *CENPV*.*L*, associated with nuclear and centrosomal processes, displays a sharply localized expression pattern, providing a clear contrast across domains [57]. Together, these genes illustrate the biological relevance of the spatial domains identified by JADE, demonstrating its ability to uncover consistent, developmentally regulated gene expression patterns across stages of brain maturation.

## 5 Conclusions

In this paper, we address the critical yet understudied challenge of joint spatial alignment and representation learning across multi-slice SRT data. We propose JADE, a unified framework that simultaneously infers spatial correspondences and learns biologically meaningful low-dimensional embeddings. Through comprehensive evaluations on human DLPFC and axolotl brain datasets, JADE demonstrates superior performance in spatial domain detection, alignment accuracy, and batch effect correction compared to state-of-the-art methods. This study provides a robust computational tool for integrating multi-slice SRT data, with the potential to advance 3D tissue reconstruction, cross-condition comparison, and spatially informed transcriptomic discovery. However, several limitations remain. First, while JADE performs well across a range of spatial transcriptomics platforms and tissue types, its performance may be affected by extreme sparsity, highly unbalanced slice resolutions, or limited overlap in tissue regions across slices. Second, JADE currently models pairwise slice alignment, which may limit its ability to fully reconstruct larger tissue volumes or resolve global correspondences across long serial sections. Third, the current framework assumes a static snapshot of spatial organization and does not account for temporal variation, which is increasingly relevant in developmental or regenerative contexts. Future work will extend JADE to handle fully joint alignment across multiple slices, incorporate spatiotemporal transcriptomics data, and integrate additional data modalities such as histological imaging or spatial epigenomics. We also aim to improve scalability for ultra-high-resolution datasets and explore methods to increase robustness under technical noise or sample heterogeneity. To the best of our knowledge, no potential negative impacts resulting from our work have been identified.

## 6 Acknowledgment

This work was supported by the National Science Foundation (NSF) and the National Institute of General Medical Sciences (NIGMS) under award number R01GM152814, by the National Institutes of Health (NIH) under award number R35GM160372, and by the NSF under award numbers DBI-2526948 and IIS-2500960.

