## Supplemental Material for "JADE: Joint Alignment and Deep Embedding for Multi-Slice Spatial Transcriptomics"

#### A Pseudocode of JADE

We summarize the training loop for JADE and its accelerated variant Fast-JADE. The notation matches Sections 2 and 3: $A_i$ is the spatial graph, $X_i$ the gene-expression matrix, $\mathcal{D}_i$ the degree matrix of $A_i$, and $D_i$ the within-slice pairwise distance matrix. JADE alternates between (i) encoding/decoding to refine embeddings and (ii) alignment in the latent space via attention and Sinkhorn normalization. Fast-JADE performs alignment at a coarser "hyperspot" level for efficiency, and then recovers full-resolution correspondences using the learned projection head.

#### B Implementation details for JADE and benchmarks

**Data Preprocessing.** We followed the standardized preprocessing workflow implemented in the SCANPY package [9] to prepare the input data for our model. For each tissue slice $i$, raw gene expression counts were normalized by library size and log-transformed. Gene expression values were then scaled to unit variance across spatial locations. The top 3,000 highly variable genes (HVGs) were identified independently in each slice, and only the intersection was retained, resulting in a shared gene set of size $p < 3000$ for both datasets (around 1500 for DLPFC and around 1000 for axolotl brain dataset). This produced two input feature matrices: $X_1 \in \mathbb{R}^{n_1 \times p}$ and $X_2 \in \mathbb{R}^{n_2 \times p}$, where $n_1$ and $n_2$ denote the number of spatial locations in slices 1 and 2, respectively.

**Spatial Graph Construction.** For each tissue slice $i$, we construct an unweighted spatial graph $G_i = (V_i, E_i)$ that captures the spatial relationships among spatial locations using spatial information $S_i$. Here, $V_i$ represents the set of spatial locations, and $E_i$ consists of edges connecting neighboring locations based on their spatial proximity. To determine connectivity between spatial locations, we compute the Euclidean distances between all pairs of locations using their spatial coordinates, then establish edges based on a defined neighborhood size $k$, where an edge is created between spatial location $i$ and spatial location $j$ if $j$ is among the $k$ nearest neighbors of $i$, thereby constructing a graph that captures the spatial relationships among spatial locations in the tissue slice. For all real data analyses, we set $k$ to be 3 The resulting graph is mathematically represented by an adjacency matrix $A_i = (a_{ij})$, where $a_{ij} = 1$ indicating a connection between spatial locations $i$ and $j$, and $a_{ij} = 0$ indicating no connection. After this step, we obtain two adjacency matrices of the spatial graph in two slices, $A_1 \in \mathbb{R}^{n_1 \times n_1}$ and $A_2 \in \mathbb{R}^{n_2 \times n_2}$.

$^{*}$Co-corresponding authors

39th Conference on Neural Information Processing Systems (NeurIPS 2025).

**Input:** Slices $(A_i, X_i)$ and distance matrices $D_i$ for $i \in \{1,2\}$; degree matrices $\mathcal{D}_i$; hyperparameters $\lambda_2, \ldots, \lambda_5$; epochs $T$; latent dim $d$; #Sinkhorn iterations $s$.
**Output:** Alignment $\Pi \in \mathbb{R}^{n_1 \times n_2}$; embeddings $H_i \in \mathbb{R}^{n_i \times d}$.
**Initialize** parameters $\{W_{ie}, b_{ie}, W_{id}, b_{id}, M, \Phi\}$
**for** *epoch* $= 1$ **to** $T$ **do**

```
// 1) Encode-Decode (GCN encoder/decoder)
```

$\tilde{A}_i \leftarrow \mathcal{D}_i^{-1/2} A_i \mathcal{D}_i^{-1/2}; \quad H_i \leftarrow \mathrm{ReLU}(\tilde{A}_i X_i W_{ie} + b_{ie});$

$\widehat{X}_i \leftarrow \mathrm{ReLU}(\tilde{A}_i H_i W_{id} + b_{id}).$

$\mathcal{L}_{\text{recon}} \leftarrow \frac{1}{n_1} \|X_1 - \widehat{X}_1\|_F^2 + \frac{1}{n_2} \|X_2 - \widehat{X}_2\|_F^2.$

```
// 2) Cross-attention & Sinkhorn for alignment
```

$S_i \leftarrow H_i M \in \mathbb{R}^{n_i \times d}$ ; `// linear projection (attention)`

$C \leftarrow \mathrm{Softmax}_{\text{rows}}\big(S_1 S_2^\top / \sqrt{d}\big) \in \mathbb{R}^{n_1 \times n_2}$

$\Pi \leftarrow \mathrm{Sinkhorn}_s(C)$ ; `// doubly-stochastic with marginals` $(\frac{1}{n_1}, \frac{1}{n_2})$

$\mathcal{L}_{\text{marginal}} \leftarrow \mathrm{KL}\Big(\frac{\Pi \mathbf{1}_{n_2}}{n_2} \,\Big\|\, \frac{\mathbf{1}_{n_1}}{n_1}\Big) + \mathrm{KL}\Big(\frac{\Pi^\top \mathbf{1}_{n_1}}{n_1} \,\Big\|\, \frac{\mathbf{1}_{n_2}}{n_2}\Big).$

```
// 3) Spatial-structure maintenance & embedding alignment
```

$\mathcal{L}_{\text{maintain}} \leftarrow \frac{1}{n_1} \|D_1 - n_2^2 \, \Pi D_2 \Pi^\top\|_F + \frac{1}{n_2} \|D_2 - n_1^2 \, \Pi^\top D_1 \Pi\|_F.$

$\mathcal{L}_{\text{align}} \leftarrow \frac{1}{n_1} \|H_1 - n_2 \Pi H_2\|_F + \frac{1}{n_2} \|H_2 - n_1 \Pi^\top H_1\|_F.$

```
// 4) Self-supervised graph contrastive loss (per slice)
```

**for** $i \in \{1,2\}$ **do**

$r_{ij} \leftarrow \frac{1}{|N(i,j)|} \sum_{k \in N(i,j)} h_{ik}$ ; `// neighbor average in the spatial graph`

Form positives $(h_{ij}, r_{ij})$ and negatives $(h'_{ij}, r'_{ij})$ by a row permutation of $H_i$

$\mathcal{L}_{\text{SCL}}^{(i)} \leftarrow -\frac{1}{n_i} \sum_{j=1}^{n_i} \big[\log \Phi(h_{ij}, r_{ij}) + \log(1 - \Phi(h'_{ij}, r'_{ij}))\big].$

**end**

$\mathcal{L}_{\text{SCL}} \leftarrow \mathcal{L}_{\text{SCL}}^{(1)} + \mathcal{L}_{\text{SCL}}^{(2)}.$

```
// 5) Update
```

$\mathcal{L}_{\text{tot}} \leftarrow \mathcal{L}_{\text{recon}} + \lambda_2 \, \mathcal{L}_{\text{SCL}} + \lambda_3 \, \mathcal{L}_{\text{maintain}} + \lambda_4 \, \mathcal{L}_{\text{align}} + \lambda_5 \, \mathcal{L}_{\text{marginal}}.$

`optimizer.step`$\big(\nabla \mathcal{L}_{\text{tot}}\big)$.

**end**
**return** $\Pi, H_1, H_2$

**Algorithm 1: JADE** (pairwise training loop)

**Evaluation metrics.** In this paper, we evaluate both alignment accuracy and representation learning quality across methods. To evaluate spatial alignment, we extend beyond conventional metrics that only consider the proportion of correctly aligned spatial location pairs sharing the same annotation (e.g., layer label) [13, 12]. While commonly used, these traditional metrics can be misleading: they often favor methods that selectively align "easy" spatial location pairs while ignoring harder regions, thereby inflating accuracy at the cost of coverage and interpretability. To address this limitation, we adopt an alignment accuracy metric that incorporates both aligned and unaligned spatial locations, offering a more robust and informative assessment. Specifically, let $\pi \in [0,1]^{n_1 \times n_2}$ be the alignment matrix where $\pi_{ij} = 1$ if spatial location $i$ in slice 1 is mapped to spatial location $j$ in slice 2 (or for soft alignment methods like JADE). We normalize each column of $\pi$ to sum up to $1/n_2$, yielding a matrix $\widetilde{\pi}$ that retains soft alignments while assigning zero mass to unaligned spots. Our alignment accuracy is then defined as $\mathrm{Acc} = \sum_{i,j} \widetilde{\pi}_{ij} \mathbf{1}(l_i = l_j)$, where $l_i$ and $l_j$ denote the biological annotations (e.g., cell type, tissue layer) for spatial location $i$ and $j$, respectively. This formulation rewards biologically meaningful alignments—i.e., mappings between spots with consistent labels—while penalizing both misalignments and omissions. When all spatial locations are perfectly aligned within the correct biological regions, $\mathrm{Acc} = 1$. Finally, to ensure symmetry, we repeat the entire procedure swapping the roles of slice 1 and slice 2 and report the average of the two scores.

Beyond alignment, we evaluate the quality of the learned representations produced by each method. We use the adjusted Rand Index (ARI) to assess clustering accuracy by comparing predicted domain labels with ground-truth annotations, providing a quantitative measure of how well the representations capture underlying biological structure. To assess batch effect removal, we compute the local inverse Simpson's index (iLISI), which quantifies the degree of slice mixing in the embedding space—higher

**Input:** Slices $(A_i, X_i)$ and distance matrices $D_i$ for $i \in \{1, 2\}$; degree matrices $\mathcal{D}_i$; hyperspot counts $m_i$; spot-to-hyperspot assignments $N^i(l)$; hyperparameters $\lambda_2, \ldots, \lambda_5$; epochs $T$; latent dim $d$; #Sinkhorn iterations $s$.
**Output:** Alignment $\Pi \in \mathbb{R}^{n_1 \times n_2}$; embeddings $H_i \in \mathbb{R}^{n_i \times d}$.
**Initialize** parameters $\{W_{ie}, b_{ie}, W_{id}, b_{id}, M, \Phi\}$
**for** *epoch* $= 1$ **to** $T$ **do**

```
// 1) Encode-Decode (same as Alg. 1)
```

$\tilde{A}_i \leftarrow \mathcal{D}_i^{-1/2} A_i \mathcal{D}_i^{-1/2}$; $\quad H_i \leftarrow \mathrm{ReLU}(\tilde{A}_i X_i W_{ie} + b_{ie})$;
$\widehat{X}_i \leftarrow \mathrm{ReLU}(\tilde{A}_i H_i W_{id} + b_{id})$.
$\mathcal{L}_{\text{recon}} \leftarrow \frac{1}{n_1} \|X_1 - \widehat{X}_1\|_F^2 + \frac{1}{n_2} \|X_2 - \widehat{X}_2\|_F^2$.

```
// 2) Hyperspot embeddings (cluster-level averaging)
```

$H_i^{\text{hyper}}[l] \leftarrow \frac{1}{|N^i(l)|} \sum_{j \in N^i(l)} h_j^i \quad \forall\, l = 1, \ldots, m_i$.

```
// 3) Hyperspot-level cross-attention & Sinkhorn
```

$S_i^{\text{hyper}} \leftarrow H_i^{\text{hyper}} M \in \mathbb{R}^{m_i \times d}$
$C^{(0)} \leftarrow \mathrm{Softmax}_{\text{rows}}\big(S_1^{\text{hyper}} (S_2^{\text{hyper}})^\top / \sqrt{d}\big)$
$\Pi^{(0)} \leftarrow \mathrm{Sinkhorn}_s\big(C^{(0)}\big)$
$\mathcal{L}_{\text{marginal}} \leftarrow \mathrm{KL}\Big(\frac{\Pi^{(0)} \mathbf{1}_{m_2}}{m_2} \,\Big\|\, \frac{\mathbf{1}_{m_1}}{m_1}\Big) + \mathrm{KL}\Big(\frac{\Pi^{(0)\top} \mathbf{1}_{m_1}}{m_1} \,\Big\|\, \frac{\mathbf{1}_{m_2}}{m_2}\Big)$.

```
// 4) Hyperspot-level maintenance & alignment losses
```

$\mathcal{L}_{\text{maintain}} \leftarrow$
$\frac{1}{m_1} \|D_1^{\text{hyper}} - m_2^2\, \Pi^{(0)} D_2^{\text{hyper}} \Pi^{(0)\top}\|_F + \frac{1}{m_2} \|D_2^{\text{hyper}} - m_1^2\, \Pi^{(0)\top} D_1^{\text{hyper}} \Pi^{(0)}\|_F$.
$\mathcal{L}_{\text{align}} \leftarrow \frac{1}{m_1} \|H_1^{\text{hyper}} - m_2 \Pi^{(0)} H_2^{\text{hyper}}\|_F + \frac{1}{m_2} \|H_2^{\text{hyper}} - m_1 \Pi^{(0)\top} H_1^{\text{hyper}}\|_F$.

```
// 5) Graph contrastive (spot-level, identical to Alg. 1)
```

Compute $\mathcal{L}_{\text{SCL}}$ as in Algorithm 1.

```
// 6) Update
```

$\mathcal{L}_{\text{tot}} \leftarrow \mathcal{L}_{\text{recon}} + \lambda_2\, \mathcal{L}_{\text{SCL}} + \lambda_3\, \mathcal{L}_{\text{maintain}} + \lambda_4\, \mathcal{L}_{\text{align}} + \lambda_5\, \mathcal{L}_{\text{marginal}}$;
`optimizer.step`$\big(\nabla \mathcal{L}_{\text{tot}}\big)$.

**end**
**Recover** $\Pi$**:** Reuse the trained $M$ to compute $C = \mathrm{Softmax}_{\text{rows}}\big((H_1 M)(H_2 M)^\top / \sqrt{d}\big)$ and then apply $\mathrm{Sinkhorn}_s$ to obtain the full-resolution $\Pi$ (as in Algorithm 1).
**return** $\Pi, H_1, H_2$

**Algorithm 2: Fast-JADE** (coarse-to-fine alignment with hyperspots)

values indicate better integration across slices. We further perform qualitative assessment using Uniform Manifold Approximation and Projection (UMAP): coloring spatial locations by their true domain labels reveals biological coherence, while coloring by slice identity visually demonstrates integration quality and batch correction performance.

**Baseline methods** We benchmarked our method, JADE, against a set of six representative baseline models, selected for their relevance to either alignment or embedding tasks in spatial transcriptomics. For alignment accuracy, we evaluated against PASTE [13], Seurat [8], and STAligner [14], three widely-used methods designed to align SRT slices based on transcriptomic similarity and spatial proximity. For evaluating the learned embeddings through spatial domain detection, we benchmarked against GraphST [6], STAGATE [2], and STAligner, all of which incorporates spatial or graph-based information to enhance the detection of spatial domains. Below we outline, for each benchmark method, the key preprocessing steps, parameter settings, and clustering or alignment workflows used in our comparisons. All pipelines begin from the same raw count matrices and spatial coordinate annotations.

- **JADE.** For each pairwise alignment task, we first selected the top 3,000 highly variable genes from each slice, then took the intersection of these two sets, yielding approximately 1,500 genes for every pair in DLPFC and 1000 genes for axolotl brain dataset. We then applied log-normalization to the expression matrix, and constructed a spot-by-gene feature matrix. For the DLPFC dataset, we normalized each inter-slice distance matrix by the minimum distance between any two distinct spots within the same slice. For the axolotl brain dataset, because the juvenile slice is slightly larger physically than the adult slice, we rescaled the coordinates of the adult slice to match that of the juvenile slice. We ended up multiplying the x coordinates of the adult slice by approximately 1.3 to eliminate any alignment bias induced by scaling. The data-driven approach of selecting $\lambda_2$-$\lambda_5$ can be found in Section E. We employed a GCN with a single hidden layer to project the original expression matrix into a 64-dimensional latent space. Following Xu et al. [11], we set the neighborhood size to $k = 3$, which has been shown to yield strong performance. Before joint training, we pretrained the model to obtain an initial estimate of the alignment matrix $\Pi$, since the embeddings $H$ are essentially uninformative at the start of optimization.

$$\mathcal{L}_{\text{pretrain}} = \mathcal{L}_{\text{SCL}} + \lambda_2 \mathcal{L}_{\text{recon}} + \lambda_3 \mathcal{L}_{\text{maintain}} + \lambda_4^0 \mathcal{L}_{\text{align}}^0 + \lambda_5 \mathcal{L}_{\text{marginal}}.$$

Similar to the Mis-Alignment loss $\mathcal{L}_{\text{align}}$, we define

$$\mathcal{L}_{\text{align}}^0 = \frac{1}{n_1} ||X_1 - n_2 \Pi X_2||_F + \frac{1}{n_2} ||X_2 - n_1 \Pi^T X_1||_F.$$

Throughout, we fixed $\lambda_4^0 = 5.0$. We used Adam optimizer with learning rate set as 0.002, number of pretraining epochs as be 200 and number of training epochs set as 800.

- **GraphST.** Following the GraphST protocol, we used the same preprocessing steps as JADE and then assembled each spot's neighborhood by integrating within-slice 2D spatial locations. Specifically, we fixed the neighborhood size to $k = 3$ spots for each query location, which is also recommended by GraphST. All other GraphST hyperparameters were left at their package defaults. After computing the spatially informed graph, we applied the mclust Gaussian mixture model for final cluster assignment, using a smoothing radius of 20 spots in the refinement stage, which again mirroring GraphST's recommended default.

- **STAligner.** We subjected the data to our standard HVG filtering and normalization pipeline before invoking STAligner's built-in neighbor selection routine. For the human DLPFC slices, we set the cutoff radius to 8 spatial units, which yielded on average approximately 5 neighbors per spot; for the axolotl brain slices, we increased the cutoff to 50 units to account for differences in spot density, also achieving roughly 5 neighbors per spot. All other STAligner hyperparameters remained at their defaults. Clustering on the aligned feature space was again performed with mclust, allowing direct comparability to GraphST and JADE.

- **STAGATE.** After the shared preprocessing steps, we turned to STAGATE's graph attention framework. We utilized its default neighbor selection within each slice, specifying a radius of 8 units for DLPFC (approximately 5 neighbor spots) and 50 units for the axolotl data (approximately 5 neighbor spots). This radius-based neighborhood ensures that each spot aggregates information from a consistent local context before passing through the graph attention layers. We retained STAGATE's default training hyperparameters and extracted the learned embeddings for clustering via mclust.

- **Seurat.** We applied the canonical Seurat integration workflow across all four consecutive slices of each sample in DLPFC dataset. We selected the top 3,000 variable genes for integration and computed 15 principal components on the combined expression matrix.

- **PASTE.** Finally, we ran the original PASTE algorithm with its default settings. Throughout, we set $\alpha = 0.1$ as recommended. No parameter tuning was performed beyond the defaults supplied by the PASTE package.

**Biological applications.** To assess the biological interpretability of the learned representations, we performed downstream analysis of domain-specific gene expression. For each spatial domain identified from the embeddings, we conducted differential expression analysis using the Wilcoxon rank-sum test to identify domain-enriched marker genes. These markers were then validated against known anatomical annotations and literature-curated gene lists. This analysis demonstrates the ability of our method to recover spatially organized, functionally coherent tissue structures and to reveal biologically meaningful gene–domain associations.

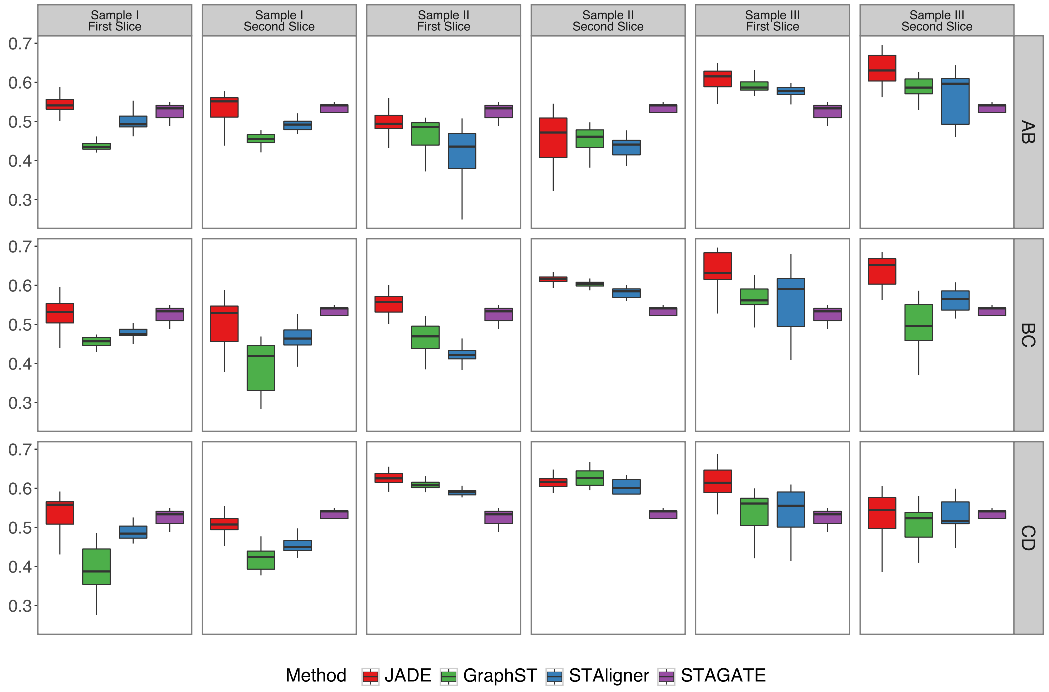


Figure 1: Adjusted Rand Index (ARI) results for three DLPFC samples (I–III, left to right), illustrating pairwise clustering of adjacent sections. Rows correspond to slice pairs AB, BC, and CD (top to bottom), and each boxplot summarizes ARI scores for both slices over 20 independent replicates.

### C Additional results for DLPFC

Complementing Figure 2(F), Figure 1 presents the ARI results for all samples and their adjacent slice pairs in the DLPFC dataset. In nearly every comparison, JADE outperforms the other methods by a substantial margin. Across all three DLPFC samples (I–III) and for each adjacent-slice pairing (AB, BC, and CD), JADE not only achieves the highest median ARI but also exhibits consistently tighter score distributions than the competing methods. For example, in Sample I's AB pair, JADE's median ARI is roughly 0.55, compared with about 0.45 for GraphST, 0.50 for STAligner, and 0.40 for STAGATE—an improvement of 0.05–0.15. Similarly, in Sample II's BC pairing, JADE scores near 0.62, while GraphST and STAligner both cluster around 0.60 and STAGATE lags at approximately 0.42. Moreover, JADE exhibits uniformly narrower interquartile ranges, particularly when compared to STAligner in Samples II and III and to STAGATE in Sample I. This indicates less variability and greater robustness to random initialization of JADE algorithm. Together, Figure 1 and Figure 2(E) highlight the satisfactory improvement achieved by JADE and underscore that joint alignment and embedding delivers more accurate and stable clustering across consecutive tissue sections.

Complementing Figure 2(D), Figure 2 displays the top 250 spot correspondences between a fixed layer of Slice A and Slice B for Sample III. Blue lines indicate correct matches, while yellow lines denote incorrect ones. Overall, JADE delivers superior alignment quality compared to all three other benchmarks: it produces markedly smoother, less noisy mappings than STAligner or Seurat across all seven layers, and it achieves higher correspondence accuracy than PASTE with only a minimal trade-off in smoothness.

In Table 1, we report, for each spatial domain within DLPFC, the alignment accuracy achieved by the top 250 correspondence links when matching Slice A to Slice B in Sample III. Each entry shows the proportion of correctly paired spots among the first 250 alignments for that domain, providing a domain-specific assessment of alignment performance. JADE emerges as the most reliable method, achieving nearly perfect accuracy (1.00) in Layer 1 and WM and maintaining at least 0.63 accuracy in every layer. PASTE also attains perfect matches in Layer 1 and WM but shows greater fluctuation in intermediate layers (ranging from 0.584 to 0.848). STAligner also delivers competitive results in early

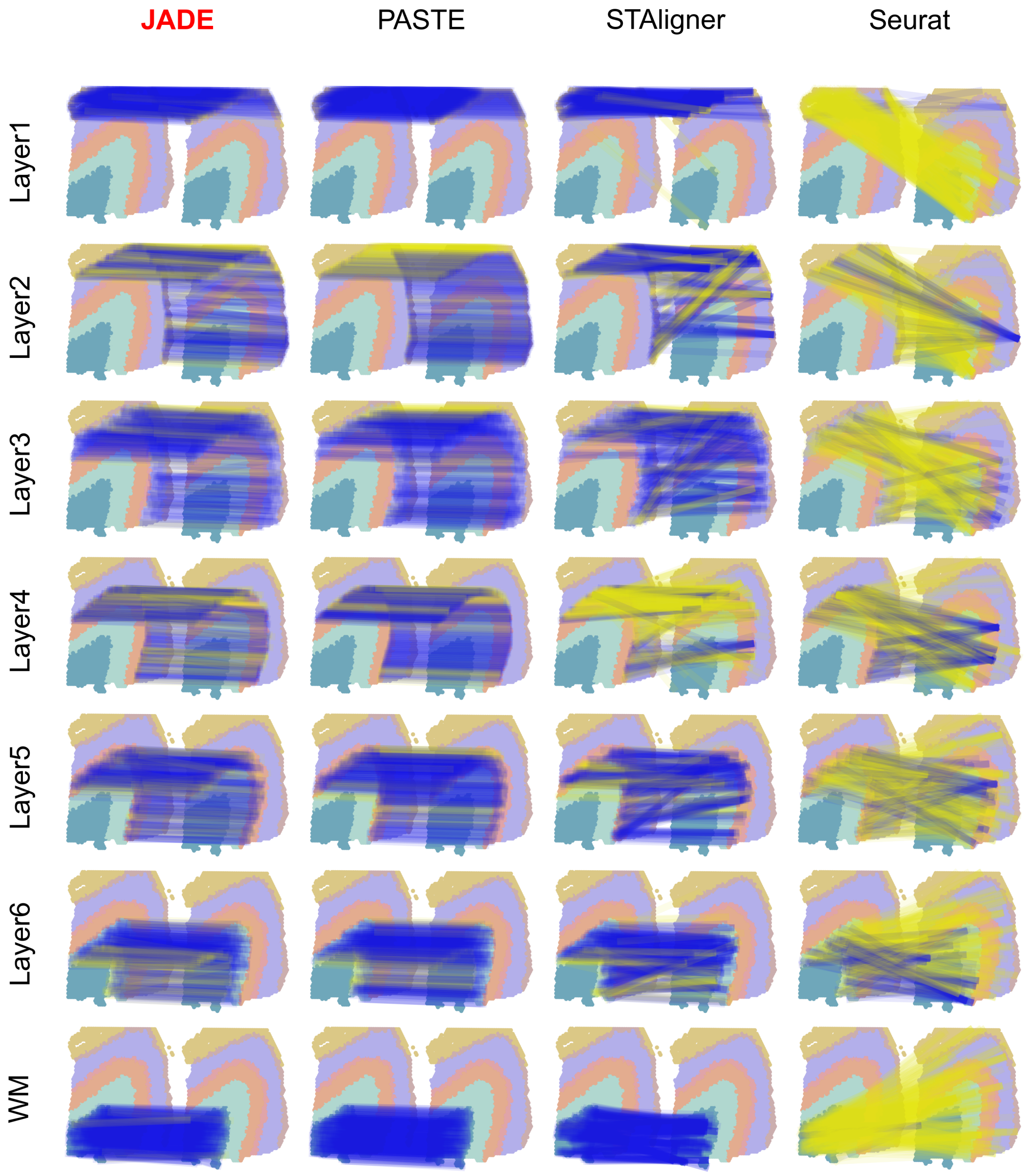


Figure 2: Illustration of alignment results from a fixed layer of Slice A to Slice B for Sample III, comparing four methods; blue lines denote correct correspondences, and yellow lines denote incorrect links.

Table 1: Accuracy of top 250 alignment correspondences by domain.

| Method | Layer1 | Layer2 | Layer3 | Layer4 | Layer5 | Layer6 | WM |
|---|---|---|---|---|---|---|---|
| JADE | 0.996 | 0.636 | **0.908** | **0.776** | **0.912** | 0.752 | **1.000** |
| PASTE[13] | **1.000** | 0.584 | 0.840 | 0.724 | 0.848 | **0.836** | **1.000** |
| STAligner[14] | 0.936 | **0.648** | 0.832 | 0.252 | 0.752 | 0.740 | 0.992 |
| Seurat[8] | 0.016 | 0.084 | 0.136 | 0.296 | 0.276 | 0.344 | 0.020 |

Table 2: Fraction of unaligned cells per domain for each alignment method.

| Method | Layer1 | Layer2 | Layer3 | Layer4 | Layer5 | Layer6 | WM |
|---|---|---|---|---|---|---|---|
| JADE | 0.000 | 0.000 | 0.000 | 0.000 | 0.000 | 0.000 | 0.000 |
| PASTE[13] | 0.000 | 0.000 | 0.000 | 0.000 | 0.000 | 0.000 | 0.000 |
| STAligner[14] | 0.739 | 0.683 | 0.709 | 0.818 | 0.738 | 0.705 | 0.790 |
| Seurat[8] | 0.953 | 0.960 | 0.932 | 0.862 | 0.889 | 0.818 | 0.984 |

layers, 0.936 in Layer 1 and 0.648 in Layer 2, but its performance drops markedly in deeper regions (as low as 0.252 in Layer 4), although it still nearly reaches perfection in WM (0.992). In contrast, Seurat fails to produce meaningful alignments across all domains, with accuracy scores below 0.35 in every layer (0.016–0.344). Table 2 reports, for each cortical layer (Layers 1–6) and white matter (WM), the fraction of spots left unaligned by each method. Both JADE and PASTE achieve a fully dense correspondence matrix, no spot remains unaligned in any domain. In contrast, STAligner fails to align a substantial fraction of spots (68–82% across layers, and 79% in WM), while Seurat leaves even more spots unmatched (93–96% in Layers 1–3, tapering to 81–82% in Layers 6 and 4–5, and 98% in WM). These results highlight that only JADE and PASTE guarantee fine-grained spot-spot alignment, whereas STAligner and Seurat produce large numbers of unaligned spots.

### D Additional results for axolotl brain dataset

Complementing Figure 3(D), Figure 3 displays the top spot correspondences between a fixed layer of juvenile slice to adult slice. The umber of correspondences is determined by half of the corresponding spots. Blue lines indicate correct matches, while gray lines denote incorrect ones. Similar to DLPFC, JADE delivers superior alignment quality compared to all three other benchmarks overall.

Table 3: Alignment accuracy of the top correspondences and average fraction of unaligned spots for each common cell type between juvenile and adult slices. For each cell type, the number of correspondences equals half of its total cells.

| Domain | # cells | JADE | PASTE[13] | STAligner[14] | Seurat[8] |
|---|---|---|---|---|---|
| CMPN | 782 | 0.357 | 0.288 | **0.448** | 0.012 |
| CP | 1500 | **0.224** | 0.157 | 0.121 | 0.069 |
| MSN | 54 | 0.685 | 0.704 | **0.741** | 0.037 |
| VLMC | 523 | 0.585 | 0.539 | **0.772** | 0.011 |
| cckIN | 99 | 0.211 | 0.169 | **0.237** | 0.019 |
| dpEX | 456 | **0.809** | 0.568 | 0.721 | 0.011 |
| mpEX | 464 | **0.623** | 0.394 | 0.591 | 0.012 |
| nptxEX | 840 | **0.751** | 0.637 | 0.744 | 0.031 |
| ribEGC | 187 | 0.337 | 0.096 | 0.690 | 0.011 |
| scgnIN | 713 | 0.421 | **0.445** | 0.372 | 0.059 |
| strpEGC | 349 | 0.630 | 0.166 | **0.745** | 0.011 |
| sstIN | 294 | 0.109 | 0.075 | **0.306** | 0.003 |
| tlNBL | 257 | 0.183 | 0.105 | **0.237** | 0.000 |
| wntEGC | 509 | **0.658** | 0.326 | 0.633 | 0.069 |
| *Avg. frac. unaligned* | — | **0.000** | **0.000** | 0.724 | 0.925 |

Table 3 compares, for each common cell type between juvenile and adult slices, the accuracy of the top correspondences and the average fraction of spots left unaligned. Across all cell types, JADE attains the highest or near–highest top-link accuracy, with values ranging from 0.105 (sstIN) up to 0.766 (nptxEX). PASTE closely follows JADE, in several cases enjoying the highest accuracy for dpEX, but generally trails by 5–10 percentage points. While STAligner exceeds JADE on individual domains (e.g. scgnIN: 0.778 vs. 0.722; sfrpEGC: 0.171 vs. 0.145), JADE remains close to STAligner in some cases (e.g. scgnIN, sfrpEGC). However, STAligner's average fraction of unaligned spots is 0.724, meaning over 70 % of cells remain unaligned. Seurat performs poorly throughout, with top-link accuracies below 0.10 in every domain and an average of 0.925 unaligned spots. Taken together, these results underscore that although STAligner can rival JADE in top-ranked matches for

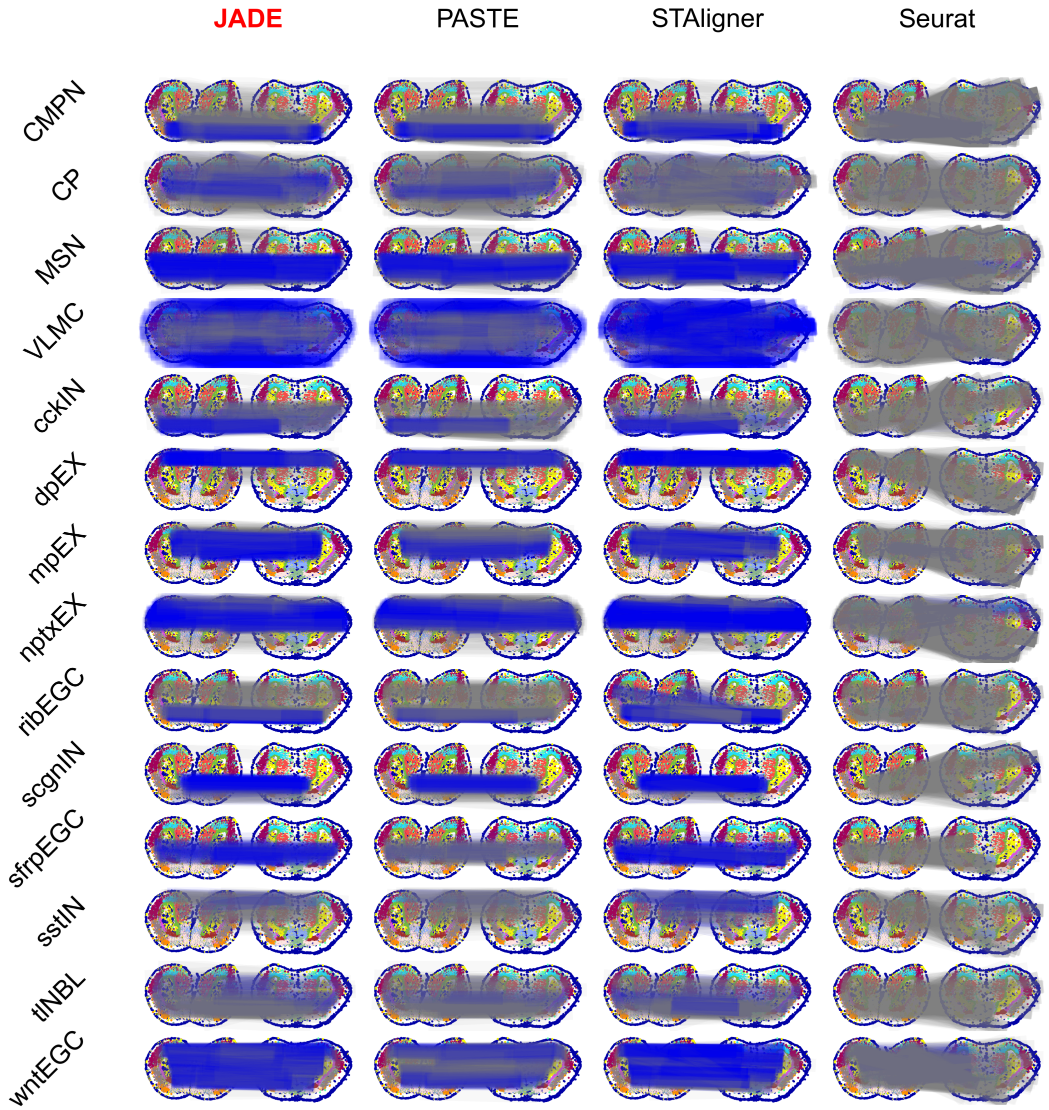


Figure 3: Alignment from a given cell type in juvenile slice to the adult slice, comparing four methods (JADE, PASTE, STAligner, and Seurat). Blue lines denote correct correspondences, while grey lines denote incorrect links.

certain cell types, its high rate of unaligned spots prevents a truly fine-grained, cell-to-cell alignment. In contrast, JADE (and PASTE) deliver both high accuracy and cell-cell alignment, ensuring no cell is left unmatched. JADE (and PASTE) combine high top-link accuracy with exhaustive alignment, leaving none of spots unaligned, making them the only methods to guarantee both precision and completeness of correspondences.

### E Hyperparameter tuning and sensitivity analysis

The training objective of JADE is

$$\underbrace{\mathcal{L}_{\text{SCL}} + \lambda_2 \mathcal{L}_{\text{recon}}}_{\text{single slice loss}} + \underbrace{\lambda_3 \mathcal{L}_{\text{maintain}} + \lambda_4 \mathcal{L}_{\text{align}}}_{\text{alignment loss}} + \lambda_5 \mathcal{L}_{\text{marginal}}.$$

The overall loss can be decomposed into three terms: the single-slice reconstruction loss, the alignment loss, and a regularization term. Throughout, we fix $\lambda_2 = 10$ and $\lambda_5 = 1$ and choose a neighborhood size of $k = 3$. The hyperparameters $\lambda_3, \lambda_4$ govern the trade-off between enforcing accurate slice-to-slice alignment and preserving meaningful, slice-specific embeddings. Within the alignment loss, $\mathcal{L}_{\text{maintain}}$ acts as a regularization term or prior belief that encodes our prior belief in the spatial coordinate similarity between the two slices. For the DLPFC dataset, we normalize each inter-slice distance matrix by the minimum distance between any two distinct spots within the same slice. We recommend using $\lambda_4 = 0.1$ and $\lambda_3 = 2.0$ by default, except in two cases—where the slices are a priori less similar—in which we set $\lambda_3 = 0.2$. To quantify slice-to-slice similarity—and thereby guide hyperparameter selection—we define the mini-max distance, which measures the average feature mismatch between corresponding spots after neighborhood smoothing. Specifically, suppose we have two slices with $n_1$ and $n_2$ spots. We first perform joint PCA and reduce the normalized gene expression matrices into low-dimensional features $Z_1 \in \mathbb{R}^{n_1 \times d}$ and $Z_2 \in \mathbb{R}^{n_2 \times d}$. Denote the adjacency matrices of these two slices by $A_1$ and $A_2$, respectively. We define the mini-max measure as follows:

$$\text{Mini-max}(Z_1, Z_2) = \frac{\text{quantile}\left\{\min_{1 \le j \le n_2} \|(A_1 Z_1)_i - (A_2 Z_2)_j\| : 1 \le i \le n_1,\ 0.99\right\}}{\sqrt{\text{mean}_i ||(A_1 Z_1)_i|| \times \text{mean}_j ||(A_2 Z_2)_j||}},$$

where the inner minimum is taken row-wise over the first slice's smoothed features, quantifying the distance between each spot of the first slice and its closest counterpart on the second slice. The mini-max distance is then defined as the 99%th-largest quantile among these minimal distances. Table 4 reports the mini–max distances for three samples across their adjacent slice pairs (AB, BC, CD). We note that the BC pairs in Sample I and Sample II have substantially larger distances than the rest. For these two cases, we set $\lambda_3 = 0.2$. Consequently, for all other cases, we set $\lambda_3 = 2.0$ to promote joint learning between slices. Figure 4 and Figure 5 present sensitivity analyses for the hyperparameters $\lambda_3$ and $\lambda_4$, respectively. Both the adjusted Rand index and the alignment score remain stable over a wide range of values: $\lambda_3$ from 0.5 to 2.0 times the default value; $\lambda_4$ from 0.05 to 0.25, deteriorating only when either hyperparameter is set to very low levels.

Table 4: Mini-max distances quantifying slice-to-slice similarity for three samples.

| Sample | AB | BC | CD |
|---|---|---|---|
| Sample I | 0.2726 | **0.5670** | 0.2609 |
| Sample II | 0.2958 | **0.4713** | 0.2764 |
| Sample III | 0.2439 | 0.2127 | 0.2368 |

Although we fixed $\lambda_2, \lambda_5$ throughout the experiment, we extend the sensitivity analysis to $\lambda_2$ and $\lambda_5$ in Table 5 to further demonstrate the flexibility of our algorithm.

### F Ablation study

Figure 6 compares JADE to versions with no mismaintain loss ($\lambda_3 = 0$) and no misalignment loss ($\lambda_4 = 0$). Results show that performance deteriorates markedly when either loss term is omitted,

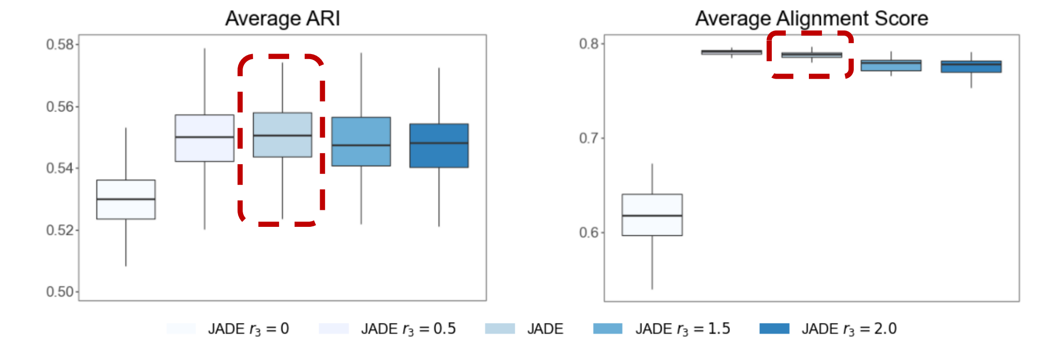


Figure 4: Sensitivity analysis: Varying $\lambda_3$, where $r_3$ is the magnifier of $\lambda_3$ against JADE. Each boxplot summarizes the outcomes of 100 independent replications.

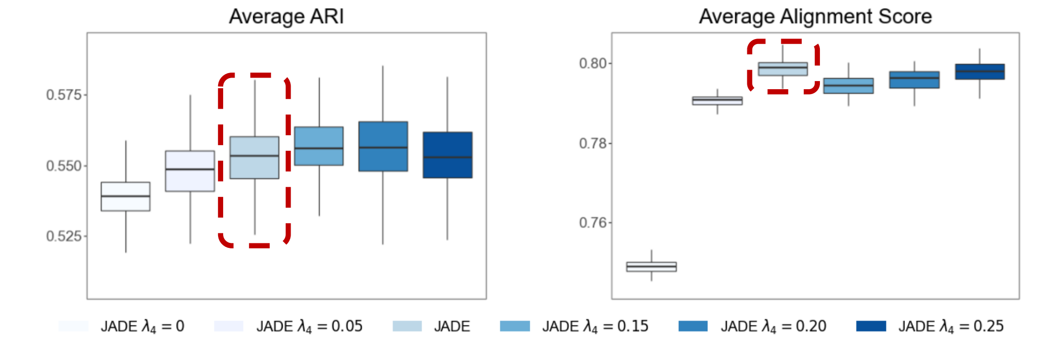


Figure 5: Sensitivity analysis: Varying $\lambda_4$. Each boxplot summarizes the outcomes of 100 independent replications.

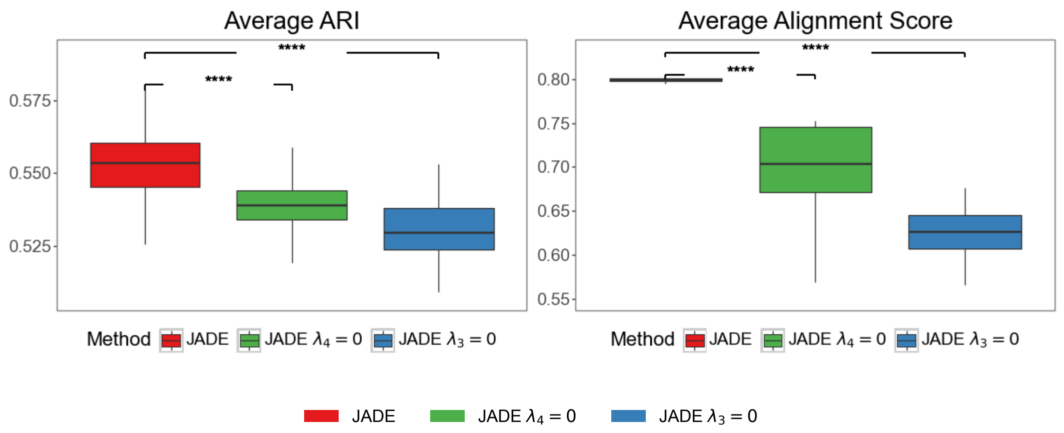


Figure 6: Ablation study: each boxplot summarizes the outcomes of 100 independent replications. We calculate the $p$-value using the Wilcoxon-rank sum test. **** stands for $p$-value lower than 0.01%

Table 5: Sensitivity analysis of $\lambda_2$ and $\lambda_5$ for JADE.

| Metric | Default | $0.5 \times \lambda_2$ | $2.0 \times \lambda_2$ | $2.5 \times \lambda_2$ | $0.5 \times \lambda_5$ | $2.0 \times \lambda_5$ | $2.5 \times \lambda_5$ |
|---|---|---|---|---|---|---|---|
| ARI | 0.550 | 0.550 | 0.564 | 0.564 | 0.547 | 0.545 | 0.552 |
| Alignment ACC | 0.788 | 0.797 | 0.790 | 0.788 | 0.801 | 0.773 | 0.787 |

demonstrating that both the misalignment and mismaintain losses are crucial for optimal embedding and alignment quality. Setting $\lambda_4 = 0$ causes the ARI to drop from 0.55 to 0.53 (a 4% decrease) and the alignment score to fall from 0.8 to 0.7 (over a 10% decrease), underscoring the critical role of the misalignment loss. Setting $\lambda_3 = 0$ causes the ARI to drop from 0.55 to 0.525 (a 5% decrease) and the alignment score to fall from 0.8 to 0.62 (about 20% decrease), underscoring the critical role of the mismaintain loss.

### G Scalability of Fast-JADE

Table 6: Runtime comparison of JADE and Fast-JADE (relative run time).

| Method | DLPFC | Axolotl Brain |
|---|---|---|
| JADE | 1.000 | 1.000 |
| Fast-JADE (with 2000 hyperspots) | 0.504 | 0.640 |
| Fast-JADE (with 1000 hyperspots) | 0.200 | 0.333 |

Table 7: Average ARI comparison of JADE and Fast-JADE in DLPFC.

| Method | DLPFC |
|---|---|
| JADE | 0.551 (0.002) |
| Fast-JADE (with 1000 hyperspots) | 0.536 (0.001) |

Table 8: GPU runtime for Fast-JADE (milliseconds per epoch).

| Number of spots per slice | 10,000 | 20,000 | 50,000 | 100,000 |
|---|---|---|---|---|
| Runtime (ms/epoch) | 14.6 | 32.3 | 84.8 | 177 |

Table 6 and 7 present the runtime and average ARI for JADE and Fast-JADE, the accelerated version of JADE using hyperspots as introduced in Section 3. We see that as the runtime reduce significantly when using Fast-JADE instead of the standard version of JADE while the ARI remains almost the same. Table 8 presents the GPU runtime per epoch for Fast-JADE. As shown in the table, Fast-JADE demonstrates approximately linear scaling with respect to the data size, maintaining efficient performance even on large-scale data.

### H Discussion

**Direct alignment against transitive alignment**. We conducted additional experiments on the DLPFC dataset (Sample III) using slices A, B, and C. Specifically, we first computed pairwise alignments $\Pi_{AB}$ (between A and B) and $\Pi_{BC}$ (between B and C) using JADE, and then derived a transitive alignment $\Pi_{AC} = \Pi_{AB} \times \Pi_{BC}$, followed by Sinkhorn normalization to enforce the doubly stochastic property. We compared this transitive $\Pi_{AC}$ with the direct alignment obtained by running JADE on slices A and C. The alignment accuracy for the direct A–C alignment was 0.767, whereas the transitive alignment via A–B–C achieved an accuracy of 0.749. These results suggest that although transitive alignment using JADE is feasible, it accumulates intermediate alignment noise, resulting in slightly lower accuracy than direct pairwise alignment. This finding highlights an important point: JADE's pairwise design supports flexible alignment across arbitrary slice pairs and enables indirect mapping when necessary, but direct alignment remains the preferred strategy due to its superior accuracy.

**Performance under unbalanced number of spots in two slices**. We conducted an experiment on the DLPFC dataset (Sample III, slices A and B), where we randomly masked a subset of spots from slice A to simulate scenarios with unequal spot counts. Table 9 reports the results obtained by varying the proportion of removed spots (10%, 25%, and 50%) in the DLPFC data.

Across all tested levels of spot removal (up to 50%), the Adjusted Rand Index (ARI) for slices A and B, overall alignment accuracy (ACC), and batch-correction metric (iLISI) remain highly stable, showing only minimal fluctuations. This robustness demonstrates that JADE effectively handles situations with substantially different spot counts between slices, making it practical for real-world applications where tissue sections vary in size.

Table 9: Performance under unequal number of spots on the DLPFC dataset.

| % Spots Removed | ARI (Slice A) | ARI (Slice B) | ACC | iLISI |
|---|---|---|---|---|
| 0% | 0.62 | 0.65 | 0.83 | 1.98 |
| 10% | 0.60 | 0.66 | 0.81 | 1.97 |
| 25% | 0.61 | 0.65 | 0.82 | 1.98 |
| 50% | 0.62 | 0.67 | 0.82 | 1.98 |

**Comparison with image-based alignment methods**. To evaluate JADE against image-based alignment methods, we conducted an experiment with GPSA [5], the only method in the benchmarking study by Hu et al. [3] that integrates both gene expression and histology features. As shown in Table 10, JADE consistently outperforms GPSA in alignment accuracy across all slice combinations in the DLPFC dataset.

While incorporating histological images can, in principle, improve alignment, their effectiveness depends heavily on image quality, resolution, and cross-sectional consistency. In practice, H&E images may be noisy, misaligned, or inconsistently stained. Moreover, these images are often obtained from adjacent rather than identical sections, limiting their spatial correspondence with transcriptomic profiles. Such discrepancies can introduce spurious or noisy signals, particularly when histological structures do not clearly delineate molecular domains.

By contrast, JADE relies on gene expression and spatial location information, which are more consistently measured across spatial transcriptomics platforms. This design enables JADE to be broadly applicable to technologies such as the Stereo-seq platform (as shown in Figure 3), where histological imaging is unavailable, and to remain robust when image quality is variable or inconsistent.

Importantly, JADE's framework is modular and can be extended to incorporate image-derived information in future work. For example, histological features could be integrated by modulating the spatial graph (e.g., assigning image informed weights to graph edges) or by combining image embeddings with expression-based representations. These extensions could further enhance JADE's utility in contexts where high-quality image data are available, while preserving its core advantage of joint alignment and representation learning. Unlike alignment-only methods such as GPSA or PASTE, JADE uniquely supports both spatial alignment and shared low-dimensional embedding, enabling a broader range of downstream analyses, including clustering, visualization, and trajectory inference.

Table 10: Alignment accuracy (ACC) comparison between JADE and GPSA on the DLPFC dataset (Samples I–III; slices A–D).

| Method | Sample I | | | Sample II | | | Sample III | | | Average |
|---|---|---|---|---|---|---|---|---|---|---|
| | AB | BC | CD | AB | BC | CD | AB | BC | CD | |
| JADE | **0.76** | **0.54** | **0.81** | **0.88** | **0.76** | **0.84** | **0.83** | **0.82** | **0.79** | **0.78** |
| GPSA[5] | 0.19 | 0.20 | 0.25 | 0.42 | 0.34 | 0.28 | 0.18 | 0.17 | 0.17 | 0.24 |

**Comparison with new benchmarks**. To further evaluate the generalizability and robustness of JADE, we conducted additional experiments on two diverse and challenging datasets: the MERFISH dataset [1] and the breast cancer Visium/Xenium dataset [4], previously used in SLAT [10]. In the latter, one slice was generated using Visium (approximately 3,500 spots and over 15,000 genes) and the other using Xenium (over 140,000 spots but only about 300 genes). As shown in Table 11, JADE consistently outperforms or matches existing methods, SLAT and STAligner, in both domain detection accuracy (Adjusted Rand Index, ARI) and alignment accuracy (ACC). These results demonstrate JADE's robustness across distinct spatial transcriptomics platforms (MERFISH, Visium, Xenium) and tissue types (brain and breast).

We note an important caveat regarding the breast cancer dataset. This dataset does not provide manually curated one-to-one ground-truth correspondences between the Visium and Xenium slices.

The Visium slice includes coarse region-level labels (e.g., *immune*, *invasive*), whereas the Xenium slice contains more fine-grained cell-type annotations (e.g., *CD8*$^{+}$ *T Cells*, *Invasive Tumor*). To enable quantitative evaluation, we manually harmonized these label sets based on biological correspondence and naming conventions. Although this approximation enables consistent comparison across methods, it introduces some ambiguity into the label-based evaluation. The reported alignment accuracy (ACC) should therefore be interpreted with caution, as it may partially reflect label mismatches rather than true misalignment. A more definitive assessment would require expert-annotated alignments, which are currently unavailable for this dataset.

Table 11: Comparison between JADE (Fast-JADE with $m_1 = m_2 = 1000$), SLAT, STAligner, and SpotScape on the MERFISH and Breast Cancer datasets. Results are averaged over 20 runs.

| Method | **MERFISH**[1] | | | **Breast Cancer**[4] | | |
|---|---|---|---|---|---|---|
| | ARI-1 | ARI-2 | ACC | ARI-1 | ARI-2 | ACC |
| JADE | **0.504** | **0.538** | **0.706** | **0.433** | **0.230** | **0.336** |
| SLAT[10] | 0.224 | 0.331 | 0.386 | 0.420 | 0.197 | 0.294 |
| STAligner[14] | 0.371 | 0.487 | 0.535 | 0.359 | 0.186 | 0.089 |
| SpotScape[7] | 0.346 | 0.265 | 0.690 | 0.277 | 0.179 | 0.312 |

[7] Yunhak Oh, Junseok Lee, Yeongmin Kim, Sangwoo Seo, Namkyeong Lee, and Chanyoung Park. Global context-aware representation learning for spatially resolved transcriptomics. *arXiv preprint arXiv:2506.15698*, 2025.
